# Rhizosphere microbes and host plant genotype influence the plant metabolome and reduce insect herbivory

**DOI:** 10.1101/297556

**Authors:** Charley J. Hubbard, Baohua Li, Robby McMinn, Marcus T. Brock, Lois Maignien, Brent E. Ewers, Daniel Kliebenstein, Cynthia Weinig

**Affiliations:** Department of Botany, University of Wyoming, Laramie, WY, USA; Program in Ecology, University of Wyoming, Laramie, WY, USA; Plant Sciences, University of California, Davis, Davis, CA, USA; Marine Biological Laboratory, Josephine Bay Paul Center, Woods Hole, MA, USA; Laboratory of Microbiology of Extreme Environments, UMR 6197, Institut Européen de la Mer, Université de Bretagne Occidentale, Plouzane, France; Department of Molecular Biology, University of Wyoming, Laramie, WY, USA

## Abstract

- Rhizosphere microbes affect plant performance, including plant resistance against insect herbivores; yet, the relative influence of rhizosphere microbes *vs*. plant genotype on herbivory levels and on metabolites related to defense remains unclear.
- In *Boechera stricta*, we tested the effects of rhizosphere microbes and plant genotype on herbivore resistance, the primary metabolome, and select secondary metabolites.
- Plant populations differed significantly in the concentrations of 6 glucosinolates (GLS), secondary metabolites known to provide herbivore resistance in the Brassicaceae, and the population with lower GLS levels experienced ~60% higher levels of aphid (*Aphis spp*.) attack; no effect was observed of GLS on damage by a second herbivore, flea beetles (*Altica spp*.). Rhizosphere microbiome (intact *vs*. disrupted) had no effect on plant GLS concentrations. However, aphid number and flea beetle damage were respectively ~3-fold and 7-fold higher among plants grown in the disrupted rhizosphere microbiome treatment, and distinct (as estimated from 16s rRNA amplicon sequencing) intact native microbiomes also differed in their effects on herbivore damage. These differences may be attributable to shifts in primary metabolic pathways.
- The findings suggest that rhizosphere microbes can play a greater role than plant genotype in defense against insect herbivores, and act through mechanisms independent of plant genotype.

## Introduction

The rhizosphere microbiome has been referred to as the “extended phenome” of the plant host because of the significant effects microbes can have on plant performance (Berendsen *et al*., 2012). Among other mechanisms, rhizosphere microbes can improve fitness in heterogeneous environments by reducing damage by herbivores (Tétard□Jones *et al*., 2007; Pineda *et al*., 2010, 2017; Badri *et al*., 2013). However, it is not known how these microbes act in combination with plant genetic pathways to modulate plant defense. For instance, plant pathways regulating the production of secondary metabolites greatly influence herbivore damage in field settings (Windsor *et al*., 2005; Kerwin *et al*., 2015, 2017, Francisco *et al*., 2016a, b). Likewise, plants grown in the presence of an intact rhizosphere microbiome experience reduced herbivory in controlled settings in comparison to those grown with an experimentally disrupted microbiome, and responses of the host plant metabolome to the microbiome are one mechanism hypothesized to underlie differences in herbivory (Badri *et al*., 2013). Given the pervasive negative effects of herbivores on plant fitness in natural settings and on crop yield (Mitchell *et al*., 2016), it would be valuable to characterize if and by what mechanisms microbes reduce herbivore damage in agroecologically relevant settings, and in particular if microbes and plant genotypes affect herbivory by similar or different metabolic mechanisms (Heil, 2008; War *et al*., 2012; Pineda *et al*., 2017).

Plant genetic variation is known to modulate plant resistance to insect herbivores by controlling the accumulation of specific primary and secondary metabolites (Kliebenstein, 2004; Bolton, 2009; Chan *et al*., 2010; Zhou *et al*., 2015; Wagner & Mitchell-Olds, 2017). Some primary metabolites are associated with cell wall composition and thickness and can thereby affect insect feeding rates (Kärkönen *et al*., 2005; War *et al*., 2012; Malinovsky *et al*., 2014). Likewise, the concentration in leaves of the primary metabolite, ascorbic acid, is directly associated with insect feeding behavior (Goggin *et al*., 2010; Zebelo & Maffei, 2015). Glucosinolates, secondary metabolites found in the Capparales, combine with myrosinases when plant tissue is disturbed, creating bioactive products that deter further feeding (Wittstock & Burow, 2010). In *Boechera stricta*, one QTL was identified that controlled both glucosinolate concentration and insect feeding (Schranz *et al*., 2009). Similarly, diverse multi-locus glucosinolate genotypes that mimic natural variants in *Arabidopsis thaliana* differed significantly in damage by chewing insects (Kerwin *et al*., 2015), clearly demonstrating that plant genotype influences the extent of damage by insect herbivores.

The microbiome also affects many aspects of plant performance including resistance to insect feeding, but it is not known if this is via similar mechanisms to those determined by natural genetic variation in plants (Berendsen *et al*., 2012; Bulgarelli *et al*., 2013; Hubbard *et al*., 2017). Microbes could influence the plant metabolome through their effects on host plant access to nutrients (Wetzel *et al*., 2016; Gomez Casati, 2016). The profile of cell wall lignin, an important physical barrier against herbivore feeding, can be affected by plant access to limiting nutrients like nitrogen, where plants with greater access to nitrogen have stronger cell walls and experience less feeding damage (Blodgett *et al*., 2005; Barros *et al*., 2015). Likewise, nitrogen and phosphorus availability can influence glucosinolate content, which can alter plant response to insect feeding (Ernst, 1998; Del Carmen Martínez-Ballesta *et al*., 2013). Rhizosphere microbes are known to improve host plant access to limiting nutrients like nitrogen and phosphorus, and could thereby indirectly influence physical and chemical plant defenses against insect herbivores (Richardson *et al*., 2009; Richardson & Simpson, 2011; Bulgarelli *et al*., 2013). Nevertheless, the magnitude of plant genotype *vs*. microbial effects on field herbivory remains unknown as does the extent to which diverse primary and secondary metabolites are influenced by rhizosphere microbes.

In the current study, we test if rhizosphere microbes and host plant population had similar (or different) effects on primary metabolites and glucosinolate secondary metabolites, and ultimately on plant responses to insect herbivores in complex field settings. We first characterize microbial community structure to test if microbiome treatments yield distinct rhizosphere microbial communities and if intact native microbiomes differ in community composition, and to evaluate what microbes may be differentially abundant across treatments. Next, we address how the rhizosphere microbiome and plant population influence the concentration of primary metabolites and glucosinolates, and how differences in the plant metabolome attributable to rhizosphere microbes and to plant population relate to damage by insect herbivores.

## Materials and Methods

### Plant Material and Growth Conditions

To investigate the influence rhizosphere microbes and host plant genotype have on the metabolome and response to herbivory, we performed 3 related types of experiments: 1) we characterized differences in rhizosphere microbial communities across distinct inoculation treatments as well as differences in GLS and primary metabolites among plant populations and between rhizosphere microbiome treatments, 2) in two independent greenhouse experiments, we determined the effect of population *vs*. rhizosphere microbiome on the extent of damage by aphids, and 3) in a field experiment, we tested the effect of population *vs*. rhizosphere microbiome on damage by flea beetles.

We used seeds from five geographically separated (minimum distance between populations = 4 km) of *Boechera stricta* populations found within the Medicine Bow, Sherman, and Sierra Madre mountain ranges of Wyoming: Crow Creek (Sherman), Road 234 (Medicine Bow), Sandstone (Sierra Madre), South Brush Creek (Medicine Bow) and Webb Springs (Sierra Madre). We collected soils from 3 of these sites to be used as microbial inoculate: Crow Creek, Road 234, and Webb Springs (**Table S1**). All seeds and soils were collected under permits from the National Forest Service (LAR1082).

For all experiments, seeds were surface sterilized using a 15% bleach solution and placed on petri plates to germinate. At the emergence of root radicles, plants were transplanted to 5cm diameter pots containing a mixture of sterilized Redi-Earth potting mix (Sungro, Agawam, MA, USA) and soil inoculate. Soil inoculate was created by mixing 30g of fresh soil with 270ml of RO H_2_O and filtered through 1,000µm, 500µm, and 212µm sieves to remove large soil particulates (van de Voorde *et al*., 2012). For the “intact microbiome” treatments, 2ml of the resulting solution was added to pots prior to planting. For the “disrupted microbiome” treatment, we filtered soil inoculate through a 0.2µm mesh to remove all microorganisms. This approach removes microbes while retaining soil nutrients, thus controlling for the potentially confounding effect of nutrients on plant performance. All experiments were performed at the Williams Conservatory at the University of Wyoming and field sites at the Agriculture Experiment Station in Laramie, WY, USA.

### Characterization of microbial communities

We harvested rhizospheres from three replicate plants grown in each of the intact (Crow Creek, Road 234) and disrupted microbiome treatments. To extract rhizosphere bacterial DNA, plant roots were agitated for 15 minutes in phosphate-buffered saline (PBS) to separate soil particles from plant roots. After removing plant roots, the soil and PBS solution was centrifuged at 3000 rcf for 15 minutes as described in Hubbard *et al*. (2017). Next, the supernatant was discarded, and 250 mg of the pellet was transferred to bead tubes from the Mobio Power Soil DNA Isolation Kit (Mobio Laboratories, Carlsbad, CA, USA). DNA was extracted following the manufacturer’s instructions. A soilless blank was included with each round of extraction to serve as a negative control. After each round of extractions, we performed PCR to quantify DNA yields and ensure reagent sterility.

Amplicon library preparation of the V4V5 region of the 16s ribosomal subunit gene (518F and 926R) and amplicon sequencing on the Illumina MiSeq platform (Illumina, San Diego, CA, USA) was performed at the Marine Biological Laboratories (Woods Hole, MA, USA) as described in Newton *et al* (2015). We used the R package *dada2* to filter and trim reads based on quality, infer error rates, merge paired end reads, remove chimeras, and assign taxonomy using the Silva reference database (ver. 128) (R Core Team, 2013; Quast *et al*., 2013; Callahan *et al*., 2016). After normalizing to account for differences between samples in read number, we performed *adonis* (permutational multivariate analysis of variance using dissimilarity matrices) and Principal Coordinate Analysis (PCoA) on Jaccard (presence-absence) and Bray-Curtis (abundance) dissimilarities based on non-rarefied reads to test if the microbial treatments were compositionally distinct using the R packages *Phyloseq* and *Vegan* (McMurdie & Holmes, 2013; Oksanen, 2015). Additionally, we used *DESeq2* to identify taxa that were differentially abundant, and may explain differential effects of intact *vs*. disrupted rhizosphere microbiomes on the plant metabolome and herbivore defense (McMurdie *et al*., 2014; Love *et al*., 2014). All sequences have been deposited into the Short Read Archive (SRA) under project number PRJNA449164.

### Characterization of the primary and secondary metabolome

To characterize the influence of plant genotype and rhizosphere microbiome on *B. stricta*’s metabolome, replicates of the Road 234 and Crow Creek populations grown in the Crow Creek, Road 234, and disrupted microbiome treatments were harvested and dried for 7 days at 65°C (*N* = 90). From each sample, 20mg of dried leaf tissue was sent to the West Coast Metabolomics Center (Davis, CA, USA), and samples were used for either high throughput glucosinolate or primary metabolite characterization (Kliebenstein *et al*., 2001a,b,c; Fiehn, 2016). Methylethyl and methylpropyl GLS were identified by comparison to previously reported profiles and relative retention times. Unidentified glucosinolates are named by their retention time (Schranz *et al*., 2009).

For analysis of the glucosinolate data, we first divided the area under peaks by dry leaf mass to obtain a measurement of glucosinolate concentration. We used Two-Way ANOVAs to characterize the effects plant population and microbiome status (intact *vs*. disrupted) had on glucosinolate concentration; we used False Discovery Rate corrections to correct for multiple comparisons. Tukey’s Honest Significant Differences *post hoc* test was used to determine the source of significant microbiome differences when observed.

For the primary metabolite data, we first corrected for drift in machine sensitivity and used Two-Way ANOVAs and False Discovery Rate corrections to correct for multiple comparisons in MetaboAnalyst 3.0 to test for the effect of plant population and microbiome status on metabolite concentration (Xia *et al*., 2016). Next, we used hypergeometric tests in MetaboAnalyst 3.0 using the *Arabidopsis thaliana* pathway library (KEGG, http://www.genome.jp/kegg/) to identify pathways significantly affected by plant population and rhizosphere microbiome. It was necessary to have separate replicates for metabolomic evaluation and tests of herbivore damage due to the comparatively small plant size and large mass of tissue needed for metabolomics.

### Plant response to insect herbivores

#### Aphids (Aphis spp.)

On August 11 2015, we planted 20 replicates of the Crow Creek and Road 234 populations in Crow Creek, Road 234, and disrupted microbiome treatments (*n* = 120), using a fully randomized design. After 6 weeks of growth, there was an unplanned infestation of aphids in the greenhouse. Aphids on the plant were counted to characterize the association of herbivory with both plant population and microbiome status. Because the original experiment was not designed to test for aphid resistance, we re-tested the effect of rhizosphere microbiome by exposing replicates of Road 234 plants in two microbiome treatments, Road 234 and disrupted, to aphids (N=18). Specifically, seedlings were transplanted from petri plates to soil on September 12 2016 and grown in the William’s Conservatory until October 20 before being placed in a fully randomized array in a greenhouse containing *Zea mays* with aphids. Aphid counts were taken daily for 10 days.

#### Flea beetles (Altica spp.)

In a fully randomized design, we grew up 20 replicates of each of the Crow Creek, Road 234, Sandstone, South Brush Creek, and Webb Springs populations in the Crow Creek, Road 234, Webb Springs, and disrupted microbiome treatments (*N* = 400). Seedlings were transferred to pots on June 6 2016 and grown in a greenhouse until July 6. On July 6, 8 replicates of each population × microbiome treatment combination were transplanted into a field with a history of flea beetle occurrence (CH, CW personal observations). Plants were checked daily for flea beetles and scored qualitatively for whole plant damage on a 0-4 scale (0 = 0-20%, 1 = 21-40%, 2 = 41-60%, 3 = 61-80%, 4 = 81-100% damage). On July 18, we harvested the 2^nd^ true leaf to quantify herbivore damage. Leaves were scanned using an Epson V700 scanner (Epson, Long Beach, CA, USA), and we used ImageJ to quantify leaf damage (Schneider *et al*., 2012).

#### Statistical Analyses

We used Two-Way ANOVAs and Tukey’s Honest Significant Differences *post hoc* tests to characterize the effect of plant population and rhizosphere microbiome status on susceptibility to aphids or flea beetles (de Mendiburu, 2016). All plots were made using the package *ggplot2*(Wickham, 2009).

## Results

### Characterization of microbial communities

After processing, we retained 1 763 803 high quality reads from a total of 2 466 906 raw reads, where read count per sample ranged from 119 042 to 313 038 reads (**Table S2**). From a planned comparison within a Jaccard presence-absence analysis, the intact (Crow Creek and Road 234) treatment differed significantly from the disrupted microbiome treatment (P = 0.014; **Fig. 1a**); further, the Crow Creek microbiome differed from that of Road 234 (P = 0.043). Likewise, from a planned comparison within a Bray-Curtis abundance analysis, the intact (Crow Creek and Road 234) treatment differed significantly from the disrupted microbiome treatment (P = 0.011; **Fig. 1b**), and Crow creek differed from Road 234 (P = 0.048). Further, differential abundance analysis (DESeq2) revealed 36 genera from 14 phyla differentially abundant between the intact *vs*. disrupted microbiome treatments (**Fig. 2**). Of the 27 genera associated with the intact microbiome treatment, 15 have been previously reported as beneficial to plant performance (Preston, 2004; Khan *et al*., 2017).

**Figure 1:**
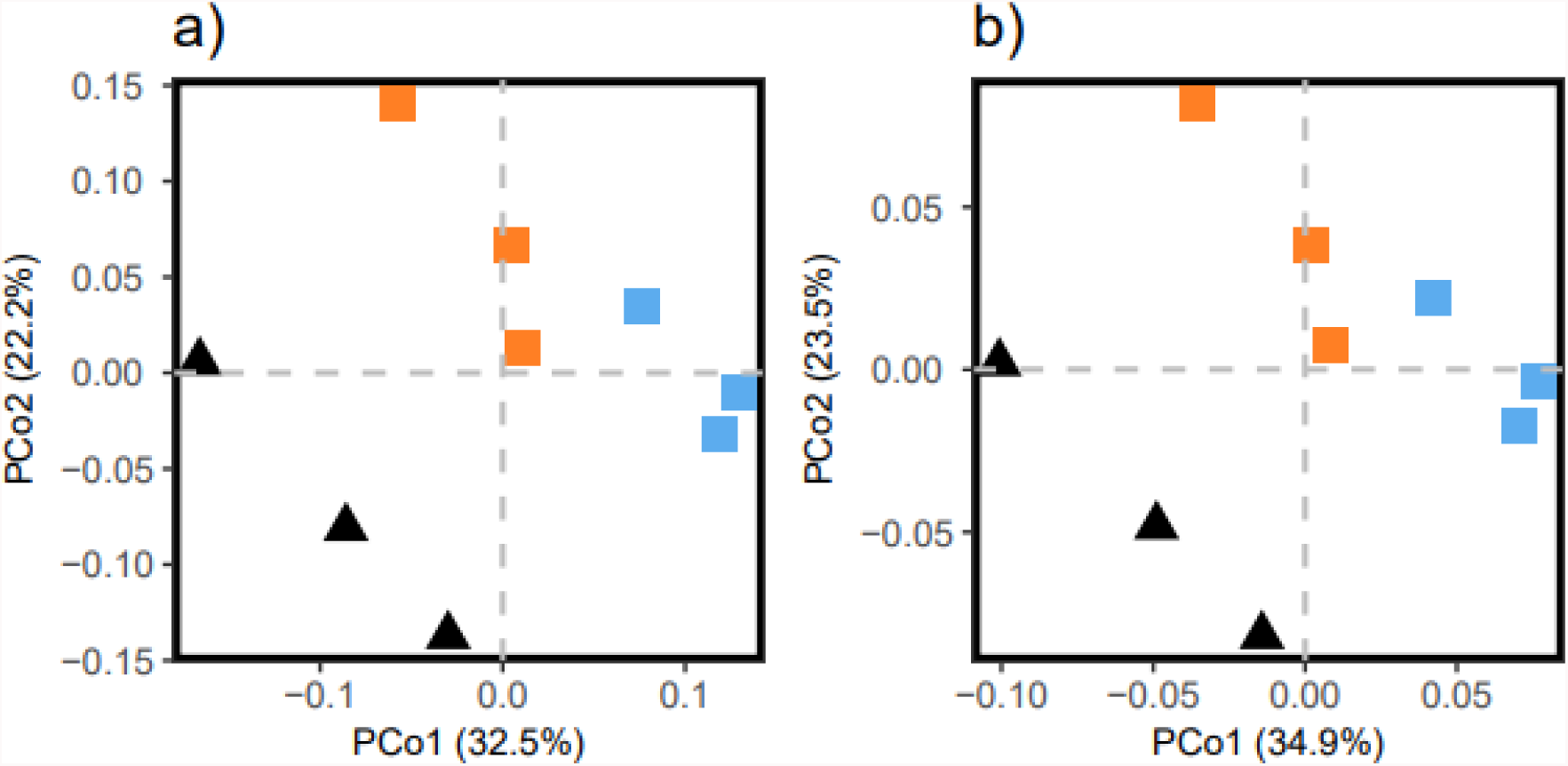
Rhizosphere communities differ between intact and disrupted microbiome treatments. Squares represent plants grown in the intact microbiome treatments and triangles represent plants grown in the disrupted microbiome treatment. Plants were grown in sterilized potting mix inoculated with Crow Creek (orange) or Road 234 (blue) microbiomes, or the disrupted microbiome (black). **a)** Principal coordinate analysis of Jaccard dissimilarities (*n* = 9). Intact and disrupted microbiome treatments differ significantly in the presence-absence of bacterial taxa (P = 0.014). **b)** Principal coordinate analysis of Bray-Curtis dissimilarities (*n* = 9). Intact and disrupted microbiome treatments differ significantly in the abundance of bacterial taxa (P = 0.011).

**Figure 2:**
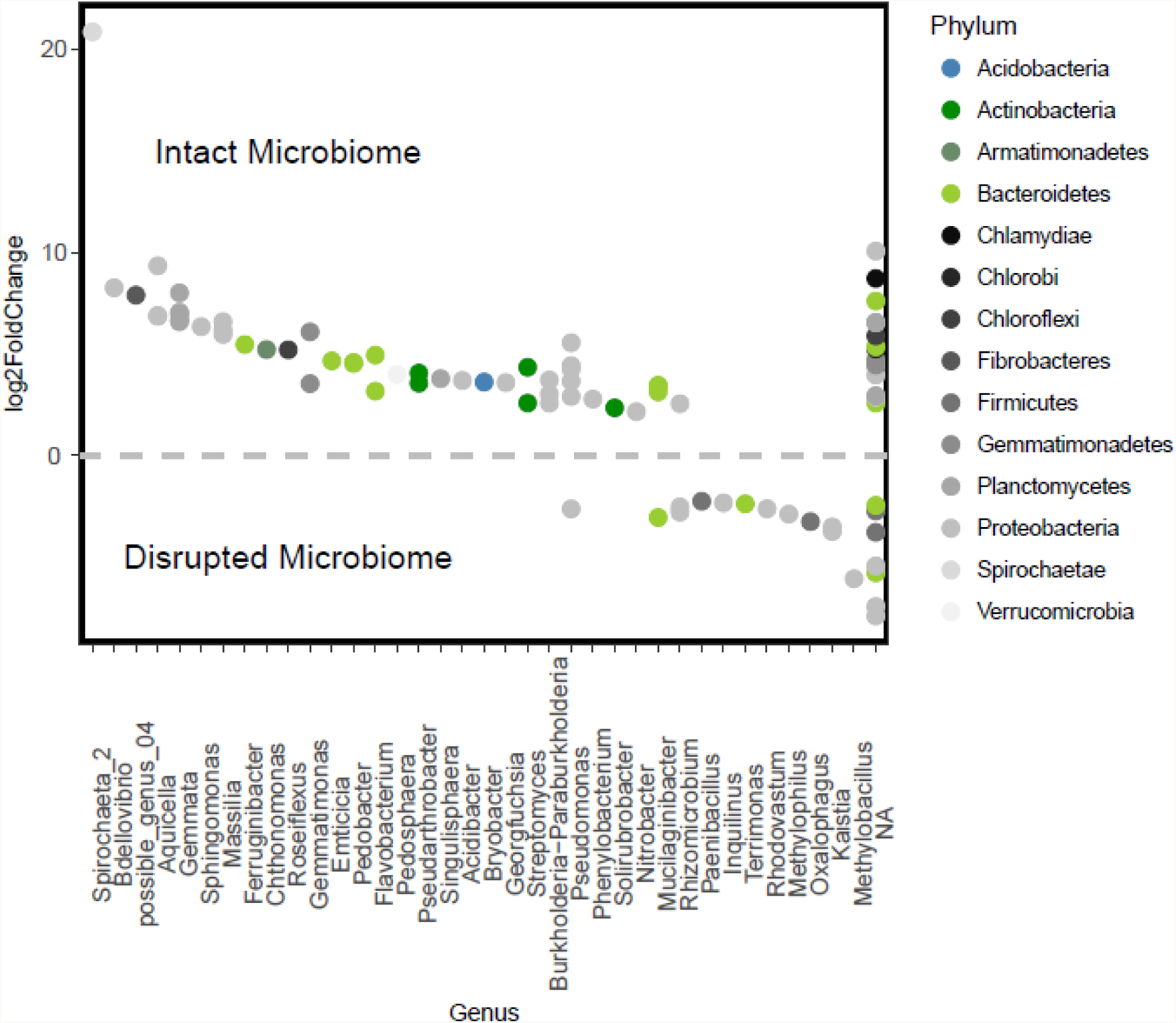
Bacterial taxa differentially abundant between intact (above the line) and disrupted (below the line) microbiome treatments.

### Characterization of primary and secondary metabolites and pathways

Plant population, rhizosphere microbiome, and their interaction significantly affected *B. stricta*’s metabolome. Six out of the nine measured glucosinolates differed significantly between populations, where five of the six were in higher concentration in plants from the Road 234 population (**Table 1**). Further, population significantly influenced the concentration of 33 primary metabolites (**Table S3**) corresponding to three metabolic pathways, all of which were at higher concentrations in Crow Creek plants (**Table 2**).

**Table 1:**
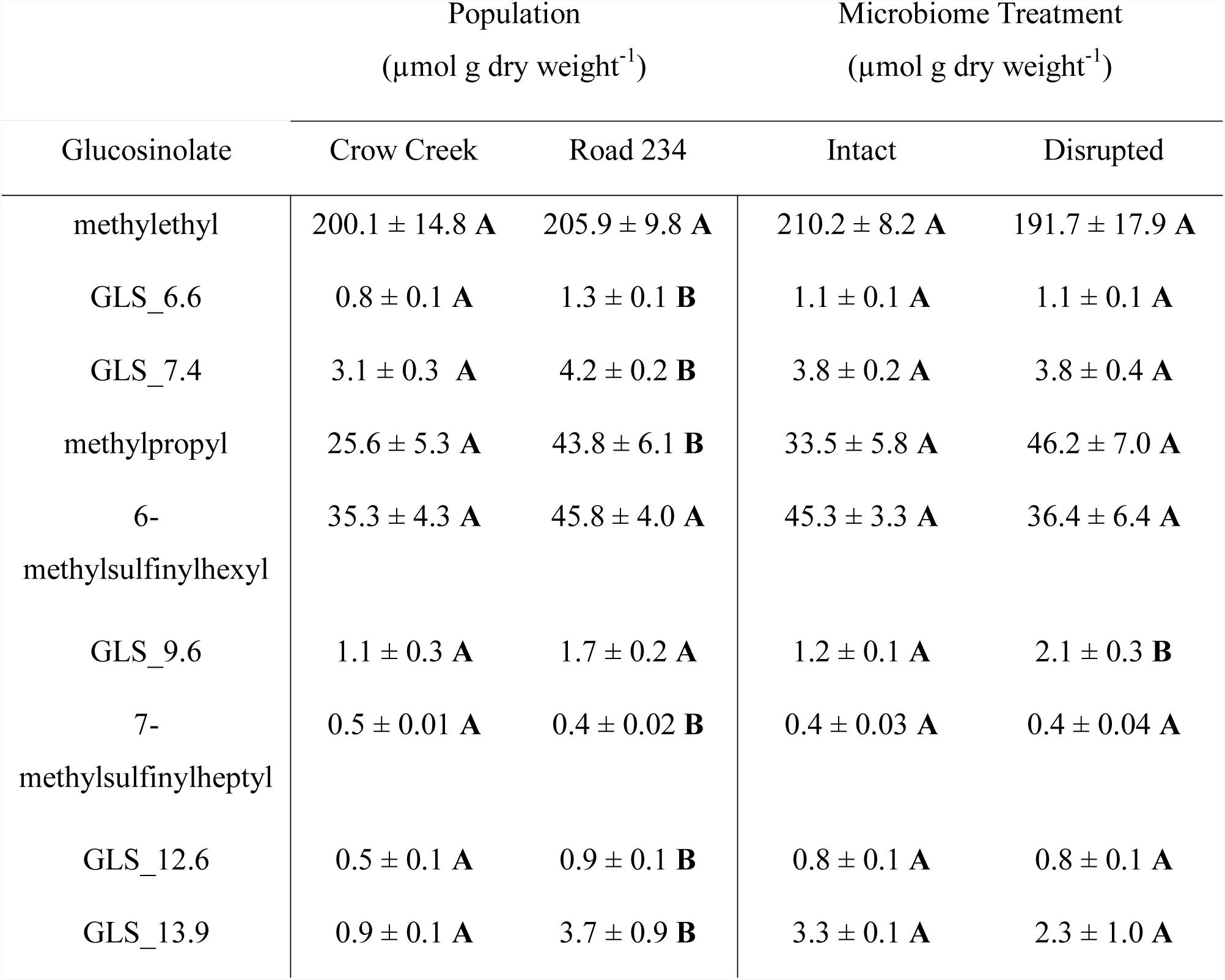
Differences in glucosinolate concentration between populations and microbiome treatments. Letters denote significant differences between population and microbiome treatments.

| Glucosinolate | Population<br>($\mu\text{mol g dry weight}^{-1}$) | | Microbiome Treatment<br>($\mu\text{mol g dry weight}^{-1}$) | |
| --- | --- | --- | --- | --- |
|  | Crow Creek | Road 234 | Intact | Disrupted |
| methylethyl | 200.1 ± 14.8 **A** | 205.9 ± 9.8 **A** | 210.2 ± 8.2 **A** | 191.7 ± 17.9 **A** |
| GLS_6.6 | 0.8 ± 0.1 **A** | 1.3 ± 0.1 **B** | 1.1 ± 0.1 **A** | 1.1 ± 0.1 **A** |
| GLS_7.4 | 3.1 ± 0.3 **A** | 4.2 ± 0.2 **B** | 3.8 ± 0.2 **A** | 3.8 ± 0.4 **A** |
| methylpropyl | 25.6 ± 5.3 **A** | 43.8 ± 6.1 **B** | 33.5 ± 5.8 **A** | 46.2 ± 7.0 **A** |
| 6-methylsulfinylhexyl | 35.3 ± 4.3 **A** | 45.8 ± 4.0 **A** | 45.3 ± 3.3 **A** | 36.4 ± 6.4 **A** |
| GLS_9.6 | 1.1 ± 0.3 **A** | 1.7 ± 0.2 **A** | 1.2 ± 0.1 **A** | 2.1 ± 0.3 **B** |
| 7-methylsulfinylheptyl | 0.5 ± 0.01 **A** | 0.4 ± 0.02 **B** | 0.4 ± 0.03 **A** | 0.4 ± 0.04 **A** |
| GLS_12.6 | 0.5 ± 0.1 **A** | 0.9 ± 0.1 **B** | 0.8 ± 0.1 **A** | 0.8 ± 0.1 **A** |
| GLS_13.9 | 0.9 ± 0.1 **A** | 3.7 ± 0.9 **B** | 3.3 ± 0.1 **A** | 2.3 ± 1.0 **A** |

**Table 2:** Population and microbiome treatment influence several primary metabolic pathways. Parentheses indicate the population or treatment in which metabolites were in the highest concentration.

| Population |  | Microbiome Treatment |  |
| --- | --- | --- | --- |
| Pathway | P – Value | Pathway | P – Value |
| Ascorbate and aldarate metabolism (**Crow Creek**) | 0.002 | Citrate cycle (**Disrupted**) | 0.002 |
| Phenylpropanoid biosynthesis (**Crow Creek**) | 0.009 | Pentose and glucuronate interconversions (**Intact**) | 0.012 |
| Glyoxylate and dicarboxylate metabolism (**Crow Creek**) | 0.041 | Glyoxylate and dicarboxylate metabolism (**Disrupted**) | 0.025 |
|  |  | Pentose phosphate pathway (**Disrupted**) | 0.027 |

In contrast to plant population, rhizosphere microbiome status influenced the concentration of only one of a total of nine glucosinolates, and for this one glucosinolate, the concentration was higher in plants from the disrupted microbiome treatment (**Table 1**).

Additionally, microbiome status affected the concentrations of 22 metabolites and four metabolic pathways (**Table S3**). Metabolites in three of the four pathways were in higher concentrations in the disrupted treatment, while metabolites in the pentose and glucoronate interconversion pathway were in higher concentrations in the intact microbiome treatment (**Table 2**). 23 metabolites were significantly affected by interactions between microbiome status and population (**Table S3**). Further, one pathway, glyoxylate and dicarboxylate metabolism, was significantly affected by both population and microbiome status, where metabolites were in higher concentrations in Crow Creek plants and the disrupted treatment (**Table 2**).

### Plant response to insect herbivores

#### Aphids

In the first experiment in fall 2015, plants from the Crow Creek population had ~3 (60%) more aphids per plant than Road 234 plants (**Fig. 3a; Table 3**). Further, in both aphid experiments, microbiome status significantly influenced the number of aphids per plant (**Table 3**). In the first experiment, plants in the disrupted microbiome treatment had ~9 (~225%) more aphids than plants in the intact microbiome treatments on average (**Fig. 3b**). Plants grown in the intact Road 234 microbiome had ~3 (113%) more aphids than those grown in the intact Crow Creek microbiome. Likewise, in the second experiment, plants in the disrupted treatment had more than 330% more aphids (~3) than plants in the intact treatment (**Fig. 3c**). Thus, the magnitude of the rhizosphere microbiome effect was larger than that of plant genotype.

**Table 3:** ANOVA table for measures of insect herbivore presence and damage.

|  | df | F value | P |
| --- | --- | --- | --- |
| **Aphid** |  |  |  |
| **Experiment 1** |  |  |  |
| *Population* | 1, 86 | 2.29 | 0.025 |
| *Microbiome Status* | 2, 86 | 26.96 | > 0.001 |
| *Interaction* | 2, 86 | 0.94 | 0.39 |
| **Experiment 2** |  |  |  |
| *Microbiome* | 1, 34 | 6.56 | 0.015 |
| **Flea Beetles** |  |  |  |
| **Leaf Damage** |  |  |  |
| *Population* | 4, 133 | 2.20 | 0.072 |
| *Microbiome Status* | 3, 133 | 28.09 | > 0.001 |
| *Interaction* | 12, 133 | 0.96 | 0.488 |
| **Whole Plant Damage** |  |  |  |
| *Population* | 4, 140 | 0.28 | 0.89 |
| *Microbiome Status* | 3, 140 | 11.76 | > 0.001 |
| *Interaction* | 12, 140 | 0.60 | 0.84 |

**Figure 3:**
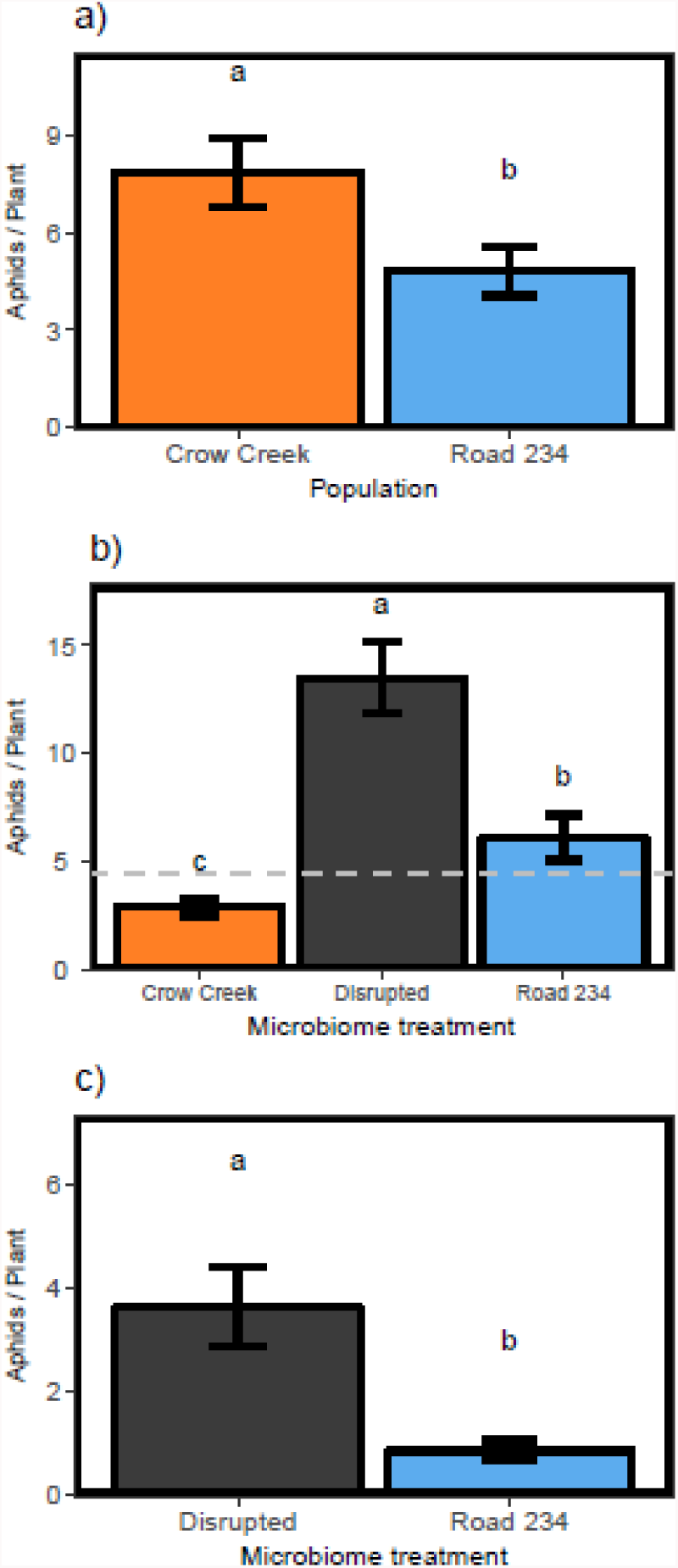
Population and microbiome treatment influences aphid prevalence. **a)** In the *fall 2015* experiment, plants from the Road 234 population (*n* = 44) had significantly fewer aphids than plants from the Crow Creek population (P = 0.027; *n* = 48). **b)** In the *fall 2015* experiment, plants grown in the disrupted microbiome treatment (*n* = 20) had significantly more aphids per plant than plants grown in the intact Crow Creek (*n* = 37) and Road 234 (*n* = 35) microbiome treatments (P < 0.001). The gray dashed line is the mean number of aphids per plant in intact microbiome treatments. **c)** In the *fall 2016* experiment, plants grown in the disrupted microbiome treatment (*n* = 35) had significantly more aphids than plants grown in the intact Road 234 (*n* = 30) microbiome treatment (P = 0.003).

#### Flea beetles

We did not observe a significant effect of plant population on leaf or whole plant damage by flea beetles (**Fig. S1; Table 3**). Plants in the disrupted microbiome treatment experienced significantly greater leaf and whole plant damage than plants in the intact treatment (**Fig. 4a, b; Table 3**). For instance, plants in the disrupted treatment, on average, experienced ~700% more leaf damage and more than 200% greater whole plant damage compared to plants in the intact treatment (**Fig. 4a, b**). The three intact microbiomes did not differ significantly in their effect on flea beetle damage (**Fig. 4a, b**).

**Figure 4:**
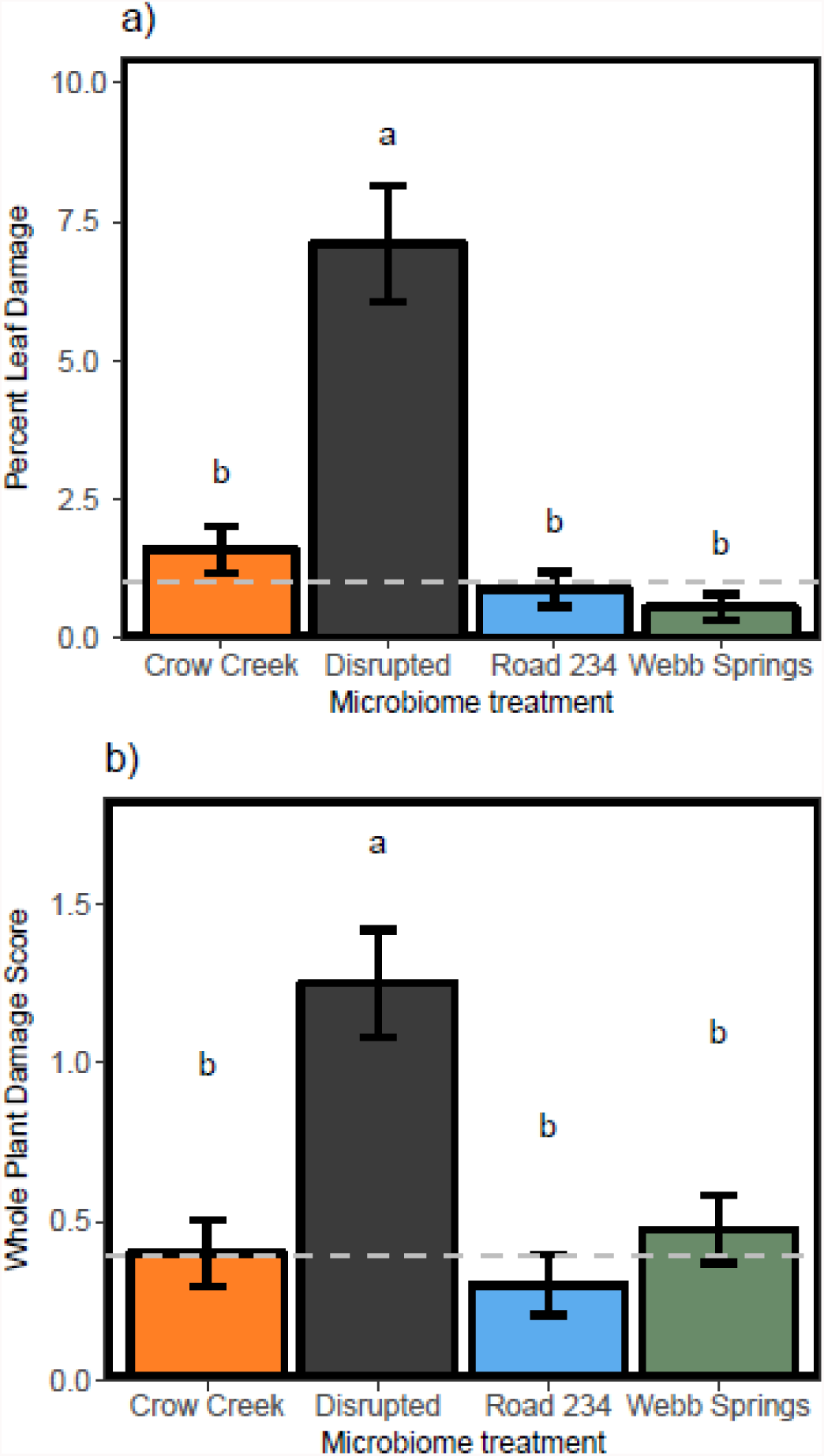
Microbiome treatment influences plant susceptibility to flea beetle damage. Gray dashed lines represent the mean damage for plants grown in the intact microbiome treatment. **a)** Plants grown in the disrupted microbiome treatment (*n* = 37) experienced significantly more leaf damage than plants grown in the Crow Creek (*n* = 38), Road 234 (*n* = 39), and Webb Springs (*n* =39) microbiome treatments (P < 0.001). **b)** Plants grown in the disrupted microbiome treatment (*n* = 40) experienced significantly more whole plant damage than plants grown in the intact Crow Creek (*n* = 40), Road 234 (*n* = 40), and Webb Springs (*n* = 40) microbiome treatments (P < 0.001).

## Discussion

The past several decades have witnessed a ground-breaking description of plant genetic controls underlying defense (Coley *et al*., 1985; Bennett *et al*., 1994; Kroymann *et al*., 2003). More recently, studies have considered the effects of the rhizosphere microbiome on plant resistance to insect herbivores (Tétard□Jones *et al*., 2007; Pineda *et al*., 2010, 2017; Badri *et al*., 2013). Although the effect of plant genotype on many metabolic defense pathways is well-characterized, the effect of microbes on the metabolome and the relative contribution of plant genotype *vs*. microbes to herbivore defense remain unclear. Here, we compared the effects of host plant population and rhizosphere microbes on metabolites and on plant responses to insect herbivores. We observed that host plant populations differed by 60% or non-significantly in susceptibility to aphids and flea beetles, respectively. Notably, intact *vs*. disrupted rhizosphere microbiome status led to 3- and 7-fold differences in defense against aphids and flea beetles, and different intact native microbiomes differed 2-fold in aphid susceptibility. Thus, the magnitude of rhizosphere microbial effect was much greater than that of plant genotype, a result consistent with the recent observation that some human phenotypes are better-predicted from the microbiome than the host genome (Sze & Schloss, 2016). The underlying mechanism of defense was seemingly distinct, because plant populations differed significantly in GLS while intact *vs*. disrupted rhizosphere microbiomes elicited differences in the primary metabolome but not GLS concentrations of the host plant.

Plant genotype has been shown to affect diverse primary and secondary metabolites linked to defense (Mauricio & Rausher, 1997; Mauricio, 1998; Kerwin *et al*., 2015; Francisco *et al*., 2016b). Here, plant population influenced several primary metabolic pathways as well as glucosinolate secondary metabolites and host plant defense against one herbivore. The three primary metabolic pathways affected by population were all in higher concentrations among plants from Crow Creek relative to Road 234 population, while 6 out of 9 measured GLS were affected by population and the majority (5) of these were found at higher concentration in plants from the Road 234 population (**Table 1, 2**). Despite the metabolomic differences between Road 234 and Crow Creek plants, we did not find significant differences in flea beetle damage between populations (**Fig. S1**). This result is consistent with previous studies of flea beetle- *Boechera stricta* interactions. In a study examining the distribution of natural *B. stricta* populations, there was an inverse relationship between *B. stricta* occurrence and flea beetle frequency despite natural variation in glucosinolate content between lines (Naithani *et al*., 2014). *B. stricta*’s inability to adapt to flea beetles may be explained by flea beetles’ ability to utilize glucosinolates as a nutrient source (Beran *et al*., 2014). By contrast, plants from the Road 234 population had significantly fewer aphids per plant than Crow Creek plants (**Fig. 3a**). It is possible that both primary metabolites and GLS contribute to reduced damage by this second insect herbivore. For example, Goggin *et al*. (2010) found that plants with higher concentrations of ascorbate were more attractive to many insect herbivores than plants with lower ascorbate concentrations, and higher concentrations of ascorbate and aldarate metabolism could correspondingly be contributing to higher rates of aphids on Crow Creek plants. Further, studies of *A. thaliana* have shown that glucosinolate concentration affects plant response to aphid feeding (Kliebenstein *et al*., 2005; de Vos *et al*., 2007; Wittstock & Burow, 2010; Zhou *et al*., 2015). Together, the metabolomic patterns are consistent with the view that Crow Creek plants may be attracting more aphids and have less protection against aphid feeding than Road 234 plants.

Our microbial treatments differed significantly in the presence-absence and abundance of bacterial taxa (**Fig. 1**), and these microbiome differences contributed could have affected defense by improving resource availability. Differential abundance analysis identified 36 bacterial genera that differed in abundance between intact *vs*. disrupted microbiome treatments (**Fig. 2**). 64 of the 84 bacterial taxa were in higher abundance in the rhizosphere of plants grown with an intact microbiome and may explain performance differences between plants across treatments. For instance, the genus *nitrobacter* was in higher abundance in the intact microbiome treatment than the disrupted microbiome treatment. In soils, *nitrobacter* is known to convert nitrite (NO_2_^−^) to nitrate (NO_3_^−^), making nitrogen more readily available to the host plant (Kumar *et al*., 1983; Richardson *et al*., 2009). As a result, plants grown in the intact microbiome treatments may have greater access to nitrogen and thus potentially higher concentrations of primary or secondary metabolites relevant to defense (Ernst, 1998; Blodgett *et al*., 2005; Del Carmen Martínez-Ballesta *et al*., 2013; Barros *et al*., 2015). Additional studies, using microbial cultures and controlled manipulations of microbial inoculates, will clarify the causal effect that different microbial taxa have on plant performance.

As in previous studies of plant-microbe interactions, we found that the disruption of rhizosphere microbiomes alters plant performance (Lau & Lennon, 2011; Badri *et al*., 2013; Chialva *et al*., 2018) and here specifically defense and several metabolites. As noted above, we found that the differences in the rhizosphere microbiome led to much greater differences in defense than did host plant population, and specifically plants grown in the disrupted microbiome treatment experienced significantly higher aphid prevalence (**Fig. 3bc**) and flea beetle damage (**Fig. 4**). While there were no differences in GLS between microbiome treatments, plants grown in the disrupted microbiome treatment had higher concentrations of primary metabolites in three pathways, and plants grown in the intact microbiome treatments had higher concentrations of metabolites in one pathway that may have altered defense (**Table 2**). Higher concentrations of metabolites in the citrate cycle among plants grown in the disrupted microbiome treatment and higher concentrations of metabolites pentose and glucuronate interconversions in plants grown in the intact microbiome treatments could contribute to the observed differences in plant response to herbivory. Upregulation of the citrate cycle is an indicator of plant stress, which can reduce plant defenses (Obata & Fernie, 2012; Bauerfeind & Fischer, 2013) and may be a factor among plants grown in the disrupted microbiome treatment. By contrast, higher concentrations of metabolites of the pentose and glucuronate interconversion pathway can increase the strength of cell walls, which could make feeding more difficult for insects (Urbanczyk-Wochniak & Fernie, 2005; Silveira *et al*., 2013) and could be important among plants in the intact microbiome treatment. It is unlikely that differences in herbivore susceptibility between the microbiomes simply reflect differences in plant vigor and poor plant growth; although plants in disrupted microbiome treatments performed poorly in response to insect herbivores in comparison to all intact microbiomes, plants from the disrupted treatment were consistently significantly larger than plants grown in the intact Crow Creek microbiome treatment in another set of experiments (CH & CW unpublished data). Together, these results suggest that plants in the disrupted microbiome treatments are more vulnerable and less well-defended than plants in the intact microbiome treatment, likely making them more susceptible and attractive to insect herbivores.

Although many metabolites and pathways were differentially influenced by either rhizosphere microbes or plant population, there was one point of overlap (**Table S3**). Metabolites in the glyoxylate and dicarboxylate metabolism pathway were in higher concentrations in plants in the disrupted treatment and among Crow Creek plants (**Table 2**). Interestingly, both experienced significantly higher rates of aphid prevalence than their counterparts. However, the role of glyoxylate and dicarboxylate metabolism in plant defense is not clear. While more work needs to be done to elucidate the role of this pathway in defense, the results suggest this is one case where plant genotype acted in a manner parallel to rhizosphere microbiome with regard to the host plant metabolome.

In sum, we observed that plant population and rhizosphere microbiome differentially affected the plant metabolome and plant response to insect herbivory in complex field settings. Plant genotype was associated with differences in glucosinolate levels as well as infestation by aphids, consistent with the well-described effects of this secondary metabolite on damage (Schranz *et al*., 2009; War *et al*., 2012; Zhou *et al*., 2015). However, the rhizosphere microbiome had much larger effects on defense than did plant genotype, and seemingly influenced defense by a different metabolomic mechanism than plant genotype, specifically a mechanism other than GLS. While we cannot exclude unmeasured metabolites, the effect of the rhizosphere microbiome may be explained by differences in primary metabolites previously implicated in plant defense or by metabolites with as yet uncharacterized effects on defense. Given the pronounced effects on herbivory, further characterizing the metabolomic intermediates between the rhizosphere microbiome and host plant defense comprises a promising avenue for future research as does functional characterization of the causal microbes.

## Acknowledgements

This work was supported by the National Science Foundation grants (IOS-1444571) to C.W., B.E.E., L.M., (IOS-1547796) to C.W., B.E.E., and D.K., EPS-1755726 to C.W and B.E.E., and a Wyoming INBRE sequencing and bioinformatics award to C.H. and C.W. We would like to thank Meredith Pratt, Ryan Pendleton, and Holly Ramseier for their help with these experiments.

## Author Contributions

C.J.H., D.K., and C.W. designed the experiments, C.J.H., B.L., R.M., and M.T.B performed the research, C.J.H, B.L., L.M., B.E.E., D.K., and C.W. analyzed the data, and all authors contributed to the writing of the manuscript.

## Supplemental Tables and Figures

**Supplemental Table 1**: Sites where seeds and soils for inoculate were collected.

**Supplemental Table 2**: Number of reads at each step of sequence data processing.

**Supplemental Table 3**: Primary metabolites significantly affected by microbiome status, population and/or their interaction.

**Supplemental Figure 1**: Populations experienced similar amounts of leaf and whole plant flea beetle damage.

